# Hard Limits and Performance Tradeoffs in a Class of Sequestration Feedback Systems

**DOI:** 10.1101/222042

**Authors:** Noah Olsman, Ania-Ariadna Baetica, Fangzhou Xiao, Yoke Peng Leong, Richard M. Murray, John C. Doyle

**Affiliations:** Department of Control and Dynamical Systems, California Institute of Technology, 1200 E. California Blvd, Pasadena, CA 91125, USA.; Department of Biochemistry and Biophysics, University of California San Francisco, 600 16th Street, Box 2542, San Francisco, CA 94158, USA.; Division of Biology and Biological Engineering, California Institute of Technology, Pasadena, CA 91125, USA.

**Author notes:** Correspondence: Noah Olsman.

## Abstract

Feedback regulation is pervasive in biology at both the organismal and cellular level. In this article, we explore the properties of a particular biomolecular feedback mechanism implemented using the sequestration binding of two molecules. Our work develops an analytic framework for understanding the hard limits, performance tradeoffs, and architectural properties of this simple model of biological feedback control. Using tools from control theory, we show that there are simple parametric relationships that determine both the stability and the performance of these systems in terms of speed, robustness, steady-state error, and leakiness. These findings yield a holistic understanding of the behavior of sequestration feedback and contribute to a more general theory of biological control systems.

## 1 Introduction

One of the central goals of systems biology is to gain insight into the design, function, and architecture of biomolecular circuits. When systems biology emerged as a field, there was a focus on the precise measurement of parameters in canonical pathways, for example those that govern glucose metabolism [1] and developmental signaling [2,3]. As both our understanding of these pathways and our quantitative measurements improved, it became apparent that many of the underlying circuit parameters are subject to large amounts of variability, despite the circuit’s overall performance being robust [4–7]. These observations led to the important insight that signaling networks have evolved sophisticated feedback control mechanisms that confer robustness, similar to those developed for classical engineering systems [8–12]. To this end, understanding the architecture and constraints of these regulatory processes is essential both to assessing the range of biological functions that they can implement and to building functional synthetic systems [13–15].

For many systems, the key to achieving robust performance is feedback control, which can provide robustness to both external noise and disturbances and to internal system variability [9,16–18]. When the system undergoes a change, such as an external disturbance or a variation in parameters, feedback can ensure that the system returns to its desired steady state with a small error [18]. Additionally, feedback control can stabilize and speedup unstable or slow processes [8,19,20]. However, feedback must be correctly designed and tuned, as it can inadvertently amplify noise and exacerbate instability [18,21]. Despite some limitations, feedback control is ubiquitous in natural biological systems, where it serves to regulate diverse processes such as body temperature, circadian rhythms, calcium dynamics, and glycolysis [8, 16, 17, 22, 23].

There are often a variety of circuit architectures capable of implementing feedback control in a biomolecular network. However, the time scale and dynamic range of their response can vary greatly depending on implementation details, such as whether the circuit relies on gene regulation [24] or post-translational modification [7]. Similarly, some circuits are robust over a broad range of inputs [25], while others may have a more modest functional range of response [4].

A particularly interesting class of biological control circuits was recently proposed by Briat et al. [26]. The authors showed that feedback implemented with a molecular sequestration mechanism is equivalent to integral feedback control [18], which guarantees perfect steady-state adaptation of the output of a network to an input signal [16]. An endogenous biological system that uses sequestration feedback to achieve perfect adaptation relies on the binding of sigma factor *σ*^70^ to anti-sigma factor Rsd [27]. Examples of synthetic biological systems that employ sequestration feedback include a concentration tracker [28, 29], two bacterial cell growth controllers [30], and a gene expression controller [31].

While integral control is a powerful tool, its stability and performance are not guaranteed to be well-behaved. Even if both the controller and the network being controlled are stable, their closed-loop dynamics may be either stable (figure 1A) or unstable (figure 1B). If the closed-loop system is stable, performance can be characterized by metrics such as tracking error, response time, leakiness, and sensitivity to disturbances. Although these metrics can be optimized individually, they can rarely all yield good results simultaneously due the constraints imposed by performance tradeoffs. These hard limits have been studied in a variety of contexts, for example in general stochastic biological control systems [32] and in the particular context of metabolic control in the yeast glycolysis system [8, 12].

Though many biomolecular circuits of interest are too complex to yield clear theoretical results that describe system-level dynamics and performance, we show in section 2.2 that a class of sequestration feedback networks can be precisely analyzed using techniques from control theory. In particular, we find that there exists an analytic stability criterion for a class of sequestration feedback systems (described in figure 1C). This stability criterion gives rise to performance tradeoffs, for example between speed and sensitivity, since fast responding controllers are intrinsically less robust. We prove these results both in the case where there is no controller degradation (section 2.3), as in the model from [26], and in the more biologically realistic context where there is such degradation (section 2.4) [33]. Though we determine many different classes of tradeoffs for the circuit, we find that they can all be viewed as different aspects of Bode’s integral theorem, which states a conservation law for the sensitivity of feedback control systems [18]. We also provide a less technical description of these results, as well as an analysis of noise in the system and simulations of synthetic circuit performance, in a companion piece [34].

These theoretical tools provide novel insight into both the analysis of endogenous biological systems and the design of synthetic systems, which we demonstrate by applying our results to a synthetic bacterial growth control circuit in section 2.5. Finally, we demonstrate in section 2.6 that it is possible to develop control architectures that will stabilize an otherwise unstable chemical reaction process. This result points towards new application domains, such as autocatalytic metabolic networks, for sequestration-based controllers that have yet to be explored in detail.

## 2 Results

Our goal here will be to develop a mathematical framework to investigate the general constraints that shape the structure of the closed-loop sequestration feedback network. For the sake of clarity we focus the results here on the simplest examples of a network regulated by sequestration feedback, however many of the results presented in this section generalize to a broader class of systems (e.g. the case with more network species and the case with controller degradation).

### 2.1 Model Description

**Figure 1:**
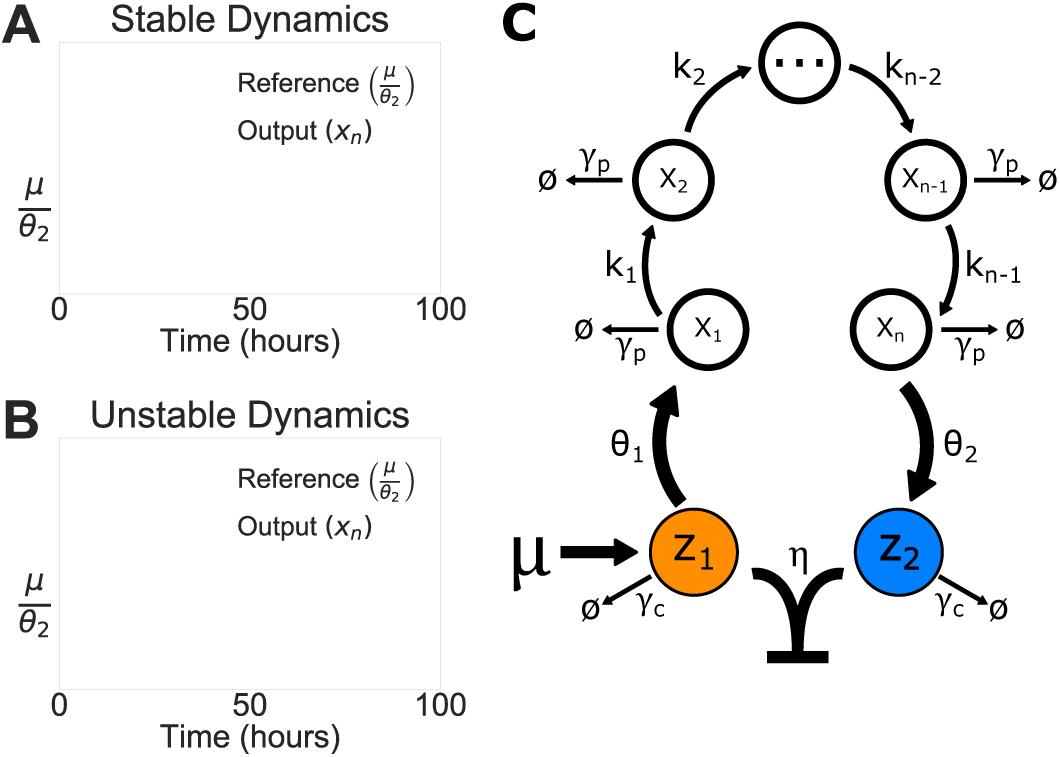
The Sequestration Feedback Network. **A)** Stable dynamics of a sequestration feedback system, where the output (solid line) precisely adapts to a reference signal (dashed line). **B)** Unstable dynamics of the same circuit, where the system is now in a parameter regime that results in sustained oscillations. **C)** A class of sequestration feedback networks. This general model has two control species, *z*_1_ and *z*_2_, and *n* process species. The two controller species are subject to a sequestration reaction with binding rate *η*. Additionally, we assume that the binding of the two controller species is much faster than their unbinding. The process species production rates are denoted as *θ*_1_, *θ*_2_, *k*_1_, …, *k*_*n*−1_. For simplicity, the process species degradation rate *γ_p_* is assumed to be equal for each *x_i_*, as is the controller species degradation rate *γ_c_*. This class of networks is defined by a simple set of possible processes where each species is only involved in the production of the next species.

We first describe the simple sequestration feedback model proposed by Briat et al. [26] with two control species (*z*_1_ and *z*_2_) and two species in the open-loop network (*x*_1_ and *x*_2_), which corresponds to the case of *n* = 2 in the general circuit diagram presented in Figure 1C with *γ_c_* = 0. In the control theory literature the network being controlled is often referred to as the process, a convention we will use in the rest of the paper.

We model the full closed-loop network using the following system of ordinary differential equations:

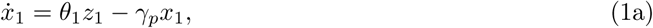

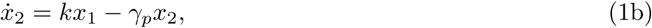

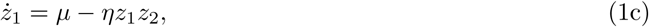

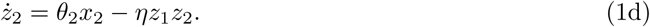

The rates *k* and *γ_p_* are production and degradation rates that are internal to the process. The parameters *θ*_1_ and *θ*_2_ are production rates that provide an interface between the network and the controller. An external reference inducer *μ* determines production rate of *z*_1_, and the two control species *z*_1_ and *z*_2_ sequester each other at the rate *η*.

While realistic models of biological circuits often have both more complex interactions and many more states, this model captures much of the important structural information about the sequestration feedback system. In particular, Briat et al. found that the network defined by (1) implements precise adaptation of *x*_2_ via integral feedback [26], as shown by the following relationship

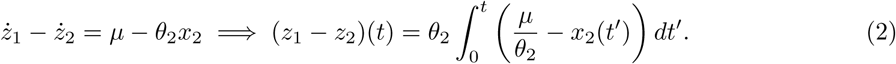

This ensures that, *if* the system is stable (i.e. ż_1_ − ż_2_ → 0), then at steady state (denoted with a *) 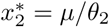. The parametric conditions that guarantee stability are not, however, obvious at first glance. Briat et al. showed general algebraic conditions that prove the existence of both stable and unstable dynamics of the linearized sequestration feedback system (using Descartes’ rule of sign), however it is not trivial to use their methods to explicitly describe stability in general.

We find that, in the limit of strong feedback (large *η*), there is a simple closed-form criterion for system-level stability. Later, we will show that a one-state network is intrinsically stable for all parameters, and that there exists a simple stability criterion for the general class of networks with many states represented in figure 1C. For the analysis, we assume both that a set of process parameters (*k* and *γ_p_*) and a desired set point (determined by *μ* and *θ*_2_) are given, and we study how stability and performance relate to the rest of the control parameters (*θ*_1_ and *η*).

### 2.2 Linear Stability Analysis

In this section, we derive an analytic criterion for the stability of sequestration feedback networks. For simplicity, we assume strong sequestration binding of the controller species (which we define mathematically later in the section).

A key difficulty in studying sequestration is the nonlinear term *η z*_1_*z*_2_ that mediates feedback in equations (1c) and (1d). Though there exist techniques to study nonlinear feedback systems, there are far more general tools available to study linear ones. While analysis of the linear system does not give guarantees about global behavior, it does allow us to characterize the local stability of the steady state to which we would like *x*_2_ to adapt. To this end, we linearize the sequestration feedback network around the non-zero steady-state value derived from equation (1):

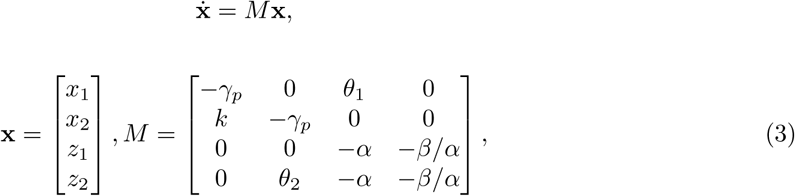

where 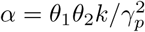 and *β* = *ημ*. We can think of *α* as representing the open-loop gain of the system, and *β* as representing the feedback gain.

In general, stability of linear systems is determined by the sign of the real parts of its eigenvalues. If they are all strictly negative, then the dynamical system is stable and the system will converge to the equilibrium point. Ideally, we would be able to directly compute the eigenvalues of *M*, however this computation corresponds to finding the roots of a fourth-order polynomial *p*(*s*) = det(*sI − M*). While this is difficult to do in general, it is possible to study stability by finding what has to be true of the parameters for the system to have a pair of purely imaginary eigenvalues, which characterizes the boundary between stable and unstable behavior. We find that, in the limit of strong sequestration (specifically *η* ≫ max(*α,γ_p_*) *·α/μ*), *M* will have purely imaginary eigenvalues *λ* = ±*iω* when *ω* = *γ_p_* = 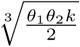. More generally, we find that the criterion for stability is

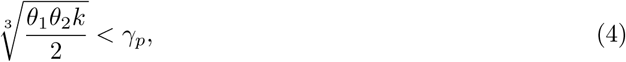

a relationship we refer to as the production-degradation inequality (proved in section S4.1). In [34], we expand on the role of *η* and how it may affect design decisions.

This implies that the closed-loop system will be stable so long as the degradation rate is larger than a constant that is proportional to the geometric mean of the production rates 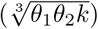. We note that, in this strong sequestration limit, inequality (4) is independent of the controller variables *μ* and *η*. Thus, this relationship tells us that stability is purely a function of the parameters describing the process and its connection to the controller, and is independent of the controller itself. Intuitively, the degradation rate sets the rate of adaptation of *x*_1_ and *x*_2_, so inequality (4) tells us that, so long as the species have a rate of adaptation that is faster than the rate of change in production, the system will be stable.

Through a more technical argument (also in section S4.1), we find that a generalized system with a chain of *n* process species has a production-degradation inequality of the form

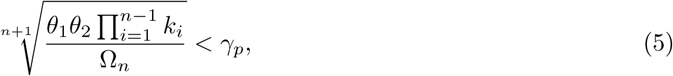

where Ω*_n_* is a constant that is only a function of the number of process species. When the system has purely imaginary eigenvalues, each species will oscillate at the frequency

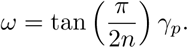

For *n* = 1 we get *ω* = tan(*π*/2) *γ_p_* = ∞, corresponding to an intrinsically stable system (i.e. it cannot oscillate or become otherwise unstable). At *n* = 2 we find *ω* = *γ_p_*, so the frequency of oscillation is equal to the process degradation rate. Since tan(*π*/(2*n*)) is a decreasing function of *n*, the frequency of oscillation will monotonically decrease as the system grows (assuming a fixed value of *γ_p_*).

Alternatively, we can interpret the parameter *α* as the open-loop gain between *z*_1_ and *z*_2_. Rearranging inequality (5), we get the inequality

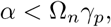

which says that the degradation rate *_p_* sets a bound on how large can be, which can be interpreted as the open-loop gain between *z*_1_ and *z*_2_, while still maintaining stability.

For simplicity, the results so far focus only on the strong feedback regime. However, we show in the supplement that there are also tractable and interesting results in the regime of weak feedback (*η* small). The results have a similar form to that of the strong feedback limit, however the direction of the inequality is reversed. The stability condition for weak feedback is:

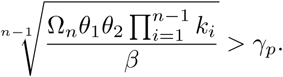

One interpretation of these results as a whole is that stability is achieved when either feedback *or* process degradation are sufficiently large, but not when both are.

##### Bode’s integral theorem and The Anatomy of a Sensitivity Function

The primary goal of any control system is to ensure that a process has a desirable response to an input signal, while minimizing the effect of external disturbances (such as noise and systematic modeling errors). While we often think of the time evolution of the full state of a dynamical system *x*(*t*), it is often useful to study the input-output relationship of a dynamical system using the (one-sided) Laplace transform

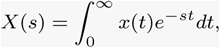

where it becomes straightforward to mathematically analyze the input-output relationship of given process.

We call functions that describe the input-output response of a system in the Laplace domain **transfer functions**, and in particular the transfer function between a reference and the output error of a system is the **sensitivity function** of a system *S*(*s*). If we take *y*(*t*) as the output state of the system (in the sequestration circuit *y*(*t*) = *x*_2_(*t*)), we denote the Laplace transform of the output *Y* (*s*). We can similarly define an input or reference signal *r*(*t*) (corresponding to *μ*) with a corresponding transformed signal *R*(*s*). We then define the error of the closed-loop system as *E*(*s*) = *R*(*s*) − *Y* (*s*), and ask how large the error of the system will be when tracking a given reference. This is given by the function

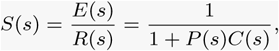

where *P* (*s*) and *C*(*s*) are the transfer functions for the open-loop process and controller, respectively.

Intuitively, when the magnitude of this function |*S*(*s*)| is small, then there will be a small tracking error between the reference signal *r*(*t*) and the output *y*(*t*). Conversely when |*S*(*s*)| is large, then there is a large tracking error. If we assume *r*(*t*) = *A* sin(*ωt*), then we can study the **frequency response** of the system |*S*(*iω*)| to a sinusoidal input with frequency *ω*.

|*S*(*iω*)| provides a way to measure system robustness, by quantifying how well a system attenuates errors to a given input. The worst-case robustness can be described by the maximum value of |*S*(*iω*)|, denoted ∥*S*∥∞. Ideally we would have |*S*(*iω*)| ≪ 1 for all *ω*. However, a deep result known as Bode’s integral theorem (proved by Hendrik Bode in 1945 [35]) states that, for an open-loop stable process, the following is true of the closed-loop response:

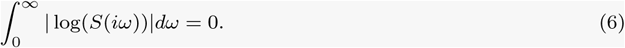

This implies that in order to reduce error in one frequency range, it *must* be increased elsewhere. This is known as the waterbed effect, and sets a fundamental limitation on the performance of any feedback control system.

**Figure 2:**
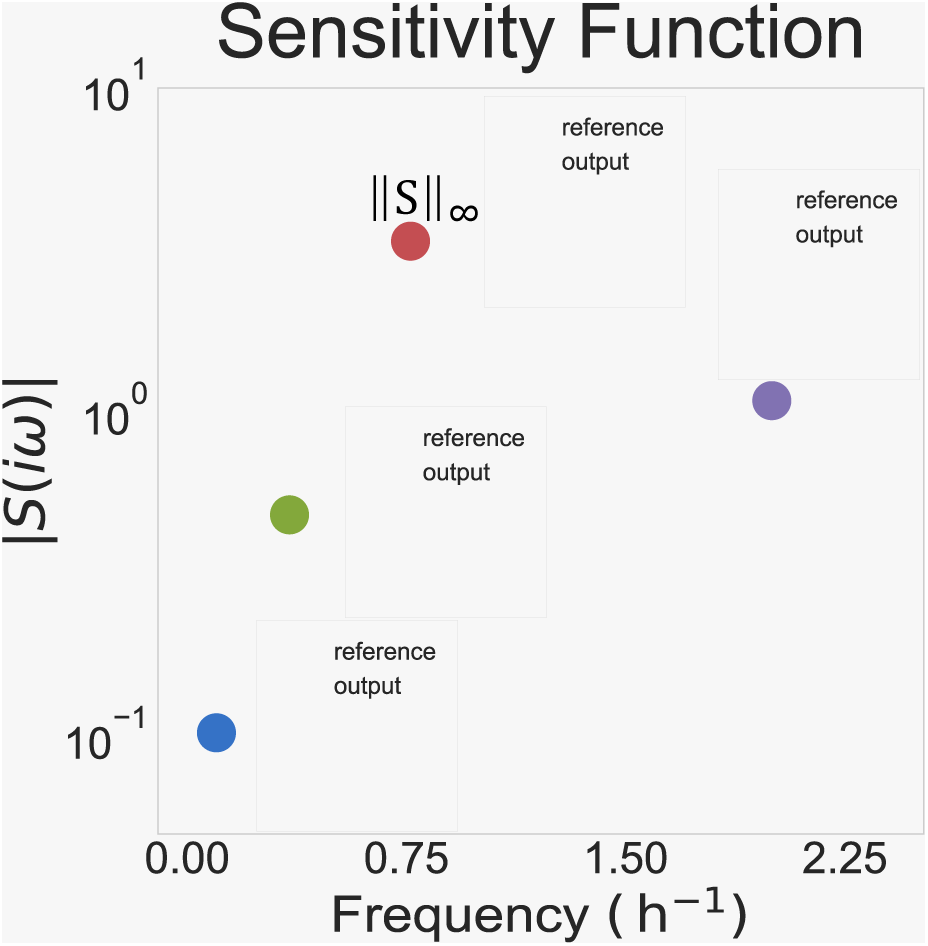
The Sensitivity Function. The sensitivity function for a system, with simulations of reference tracking dynamics for various inputs. We see that when |*S*(*iω*)| < 1, the system has small error and performs well (blue and green). At the peak |*S*(*iω*)| = ∥*S*∥∞. (red), we see that the output magnitude is not only amplified, but also phase shifted such that it is almost exactly out of sync with the reference. At high frequencies (purple), the reference is changing so quickly that the system can barely track it.

### 2.3 Performance Tradeoffs and Hard Limits

While inequality (4) gives us a binary condition that determines stability, it does not directly tell us about overall performance of the system. We know when the system becomes unstable, but it is unclear how the system behaves as it approaches instability. Let

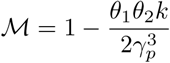

be a measure of how far the system is from going unstable that we will refer to as a *stability measure* of the system. For simplicity the analysis here will focus on the *n* = 2 species case, however the results naturally generalize for arbitrary *n*. From inequality (4), we have that ℳ = 0 implies instability, and that the larger ℳ is the further the system is from becoming unstable. Intuitively it seems that the system should become increasingly fragile to disturbances as ℳ approaches zero. Conversely, we can increase ℳ by decreasing the production rates *θ*_1_, *θ*_2_, and *k*, but this will slow down the dynamics of the system and could potentially hurt performance.

To analyze this problem, we will study the sensitivity function *S*(*s*), which is the transfer function between the reference signal and the output error of the system [18]. This transfer function captures the effect of external disturbances on the output error of a system, in this case, *x*_2_. The sensitivity function is described in greater detail in the box above.

While there are many different ways to characterize robustness, generally we consider a system to be robust if there no small change in parameters that would cause it to become unstable. A mathematically equivalent interpretation is that a system is robust when its worst-case error when tracking references (i.e. the maximum value of *S*) is small [18]. For the *n* = 2 case of the circuit in figure 1, we have (see section S4.2):

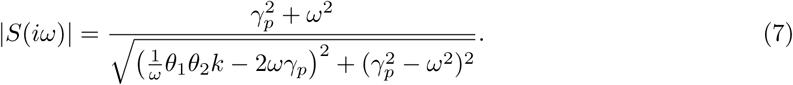

The robustness of a system can be formally quantified by ∥*S*∥_∞_. = max*_ω_*|*S*(*iω*)|, the maximum magnitude of the sensitivity function across all frequencies (in mathematics, the quantity ∥*·*∥_∞_ is referred to as the infinity norm of a function). The quantity ∥*S*∥_∞_ describes the worst-case disturbance amplification for the system to an oscillatory input. If ∥*S*∥_∞_ is in some sense small enough to be manageable, then values of |*S*| across all frequencies are also small and the system is robust to any disturbance. If ∥*S*∥_∞_ is large enough to be problematic, then there is at least one disturbance against which the system is fragile.

Directly computing ∥*S*∥_∞_ in terms of the parameters of a system is difficult in general, but it is sometimes possible to compute good lower bounds that yield insight into a system’s robustness. To this end, we find that (see Section S4.2 for a detailed proof):

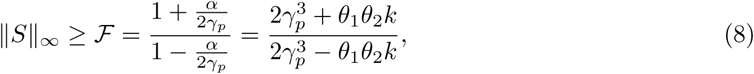

with equality when

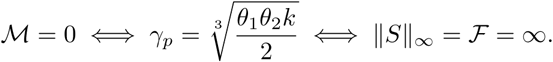

The fragility bound ℱ is constructive, in that we can write down the frequency *ω*\* that achieves it:

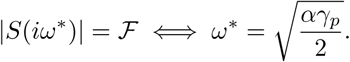

For a given constant reference *μ/θ*_2_, we use equation (8) to derive a tradeoff between fragility and response time (which we quantify with 1/*θ*_1_). Figure 3A shows this tradeoff curve for a particular set of parameters as θ_1_ varies, with the corresponding dynamics shown in figure 3B. We see from the latter plots that the response time (1/*θ*_1_) and fragility (ℱ) correspond directly to the rise times and oscillatory behavior of simulations in figure 3B. Figure 3C shows the corresponding sensitivity functions, with colored dots corresponding to values of ℱ. Here we can clearly see Bode’s integral theorem (equation (6)) at work, in that the area above and below the dashed line (corresponding to log |*S*(*iω*)| = 0) is always equal. We see that, as dynamics become more oscillatory, ∥*S*∥_∞_ becomes large.

Because we have fixed *μ/θ*_2_ and assumed that is large, the only remaining control parameter to vary is *θ*_1_, so there will only ever be one meaningful tradeoff dimension to study for this system. In the next section, we present results for the case with non-zero controller degradation rates. This model is both more biologically realistic and provides a richer tradeoff space to analyze.

**Figure 3:**
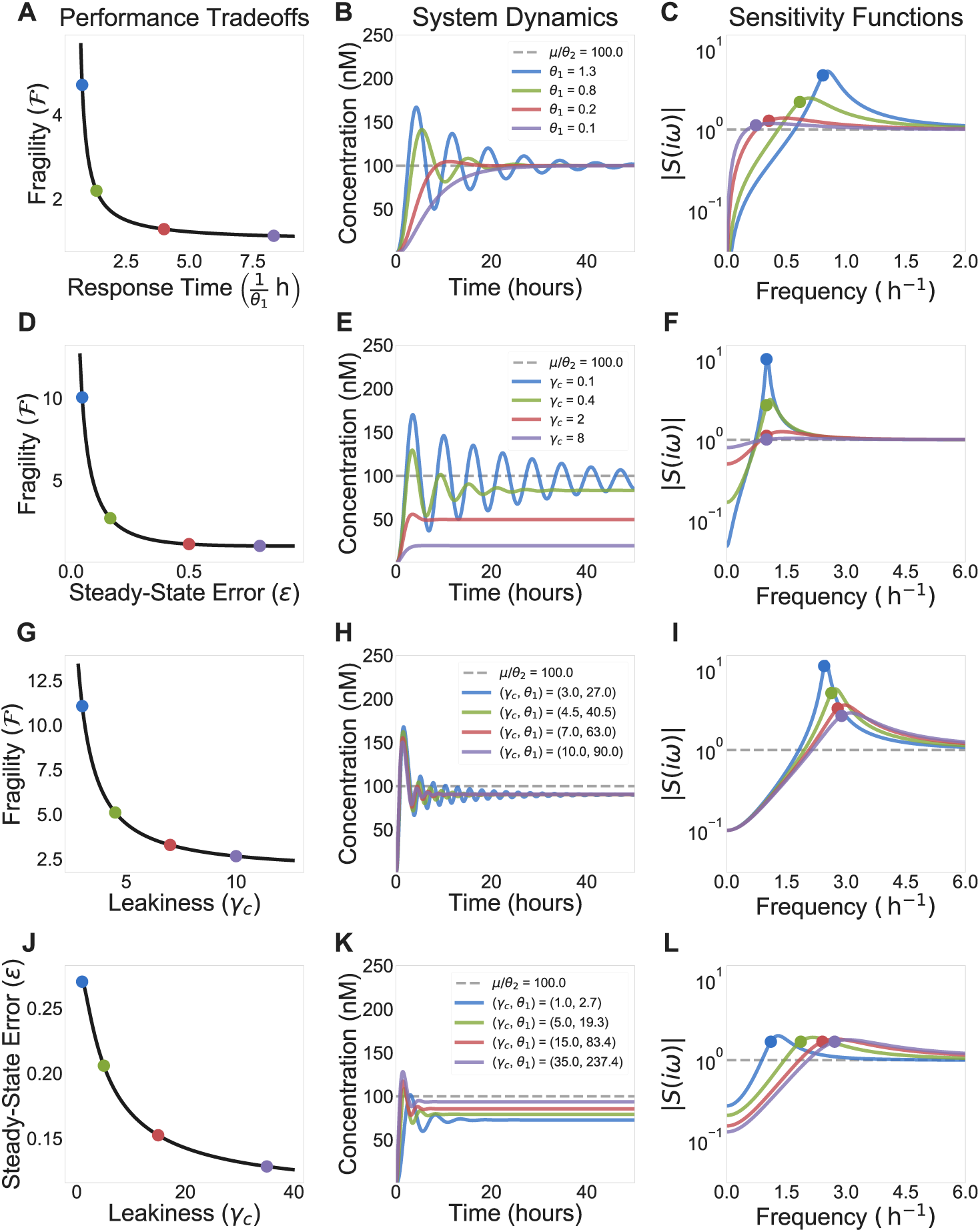
Hard Limits and Performance Tradeoffs in Sequestration Feedback Circuits. **A)** We see the relationship between speed and fragility in the sequestration feedback system. Speed can be characterized in terms of any of the production rates of the system (here we vary 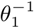), where higher production rates lead to a faster response. Fragility is defined as a lower bound on the maximum value of the sensitivity function ∥*S*∥_∞_ as defined in equation (8). **B)** Time-domain simulations corresponding to different points on the tradeoff curve in A. We see that speed and fragility naturally relate to the rise time and settling time of the system. **C)** Sensitivity functions for various parameter values. We see what is known in control theory as a waterbed effect, where better attenuation of disturbances at low frequencies necessarily implies worse amplification of disturbances at higher frequencies as a result of equation (6). The colored dots correspond to values of ℱ computed using equation (8). **D)** Here we set *γ_c_* > 0 and observe the effects of controller degradation being varied on its own. We set *θ*_1_ = 2 h^−1^ so that, if *γ_c_* = 0, the system would be unstable. We see that increasing *γ_c_* decreases fragility, at the cost of introducing steady-state error, which is illustrated in the dynamics shown in panel **E. F)** The corresponding sensitivity functions also illustrate the tradeoff, where the peak of |*S*(*iω*)| (fragility) decreases as the value of |*S*(0)| (steady-state error) increases. *F* is now computed using equation (13). **G-I)** In these plots we vary both *γ_c_* and *θ*_1_ such that *θ*_1_/*γ_c_* = 9, corresponding to *ε* = .1 in equation (11). We now observe a tradeoff between fragility and leakiness, the latter being captured by how much turnover of *z*_1_ and *z*_2_ is introduced by *_c_*. **K-L)** Finally, we can instead hold ℱ constant and numerically solve for *θ*_1_ given a value of *γ_c_*. This introduces a tradeoff between steady-state error and leakiness. In all simulations *k* = *θ*_2_ = *γ_p_* = 1 h^−1^, *η* = 1000 h^−1^ nM^−1^, *μ* = 100 nM h^−1^.

### 2.4 The Effects of Controller Species Degradation

In the previous sections, we assumed that the controller species does not degrade and we derived an analytic stability criterion for closed-loop sequestration feedback networks. Fulfilling the stability criterion ensures that the sequestration feedback network precisely adapts. As discussed, perfect adaptation is a desirable property because it facilitates disturbance rejection and robustness despite variability process dynamics. However, the literature suggests that implementing sequestration feedback with no controller species degradation is challenging [36,37]. Because of this, we will now extend our analysis of stability, performance, and tradeoffs to sequestration feedback networks with nonzero controller species degradation rates.

To model the effects of controller species degradation, we modify equations (1c) and (1d) such that,

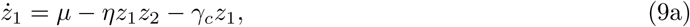

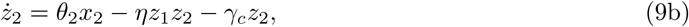

where *γ_c_* is the degradation rate of the control species *z*_1_ and *z*_2_.

Including the controller species degradation rate in the sequestration feedback network model changes its properties of stability and performance. In particular, the closed-loop sequestration feedback network has zero steady-state error for *γ_c_* = 0, whereas if *γ_c_* > 0 then there will generally be some non-zero error in *x*_2_.

In the limit of strong sequestration, we can analytically compute the steady-state values of the system species and bound its sensitivity function. While it is somewhat more complicated to compute even the steady-state values of each species for this system, we show (see section S4.3.1) that, in the limit of large *η*, it is possible to derive a simple approximate formula for 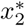:

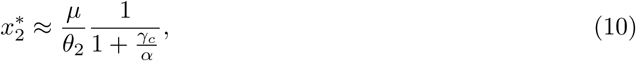

from which all other steady-state values can be derived. Under the strong feedback assumption, *x*_2_ no longer precisely adapts to the set point *μ/θ*_2_, but rather will have some amount of steady-state error determined by the ratio *γ_c_/α*. The relative error in 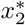 can be quantified by the relationship

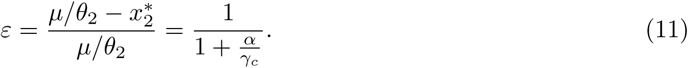

We see that *γ_c_* = 0 ⇒ *ε* = 0, corresponding to our previously results that precise adaptation is achieved when there is no controller degradation. Using this simplified expression, the relative steady-state error function can be bounded. For example, if we are interested in obtaining *ε* < .1, then we can choose a controller degradation rate such that 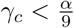.

Moreover, we can also derive the corresponding stability criterion (see section S4.3.3). Here we present the stability criterion for the two process species case:

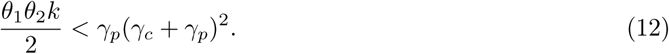

This reduces to inequality (4) when *γ_c_* = 0, and shows that *γ_c_* > 0 leads to a increased stability measure. If we only consider variations in *γ_c_*, then the combination of equation (11) and inequality (12) yields yet another tradeoff. As *γ_c_* increases, the system becomes increasingly stable at the cost of worse steady-state error (see figure 3D and E).

In section S4.3, we derive a general stability criterion that depends on comparing the controller and the process species degradation rates for *n* > 2 process species. When the process degradation rate is much larger that the controller degradation rate or the two are comparable, then the stability criterion is the same as the production-degradation inequality. However, when the process degradation rate is much smaller than the controller degradation rate, then the stability criterion relies on the controller degradation term to compensate for the slow process degradation rate. Since the process network is slow, the sequestration feedback network is challenging to stabilize and its performance can be very poor.

We now focus on analyzing the properties of the sensitivity function and the tradeoff it introduces. Figure 3F shows the corresponding sensitivity function for this system. One major difference between these sensitivity functions and those in figure 3C is that we now have |*S*(0)| > 0. This is directly related to the steady-state error in equation (10), as we can think of a signal with frequency *ω* = 0 as a constant reference. A convenient property of the sensitivity function is that |*S*(0)| = *ε*, so the previously mentioned tradeoff between robustness and steady-state error can be recast as a tradeoff between |*S*(0)| and ∥*S*∥_∞_. In figure 3C we see that log |*S*(0)| = −∞, corresponding to |*S*(0)| = 0, implying *ε* = 0 steady-state error. Because of the waterbed effect, increasing |*S*(0)| has a tendency to reduce ∥*S*∥_∞_. This can be seen directly by deriving a bound similar to the one in equation (8) for the case *γ_c_* > 0:

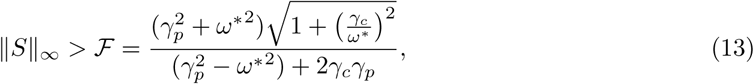

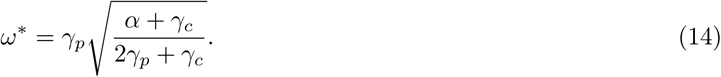

Though ℱ is now more complicated, we can see that it will scale as 𝒪(1/*γ_c_*) for small *γ_c_*. This tells us that increasingly *γ_c_* has the potential to reduce ℱ. In figure 3D we see this effect, where ℱ asymptotically decreases to 1 as *γ_c_* (and consequently *ε*) increases. It is also straightforward to check that ℱ reduces to equation (8) when *γ_c_* = 0.

So far we have shown what happens when the control parameters *θ*_1_ and *γ_c_* are varied individually, however it is also interesting to study what happens when they are varied such that a particular performance characteristic is held constant. Figure 3G, H and I demonstrate the system’s response when we vary *θ*_1_ and *γ_c_* such that the steady-state error *ε* is fixed. This sort of variation can be interpreted as changing the turnover rate, and consequently the leakiness, of the controller. This leakiness can also be thought of decreasing efficiency, as it means that control molecules are degraded before ever being involved in feedback. By increasing *γ_c_*, we make the system less efficient because the controller spends resources producing and then degrading molecules of *z*_1_ and *z*_2_. Figure 3G shows that highly efficient controllers are more fragile than less efficient ones. We can also see this in figure 3I, where the integrated area of |*S*(*iω*)| gets spread out over high frequencies, rather than having a large and narrow peak. This leads to a lower value of ∥*S*∥_∞_ and a corresponding increase in robustness. Conversely, we can fix ℱ and see how *ε* changes with leakiness. In figure 3J and K we see that highly efficient controllers have worse steady-state error, and as the controller becomes less efficient *ε* improves. This can be observed in figure 3L, Where |*S*(0)| is reduced as *γ_c_* increases. Because |*S*(0)| is decreasing and ∥*S*∥_∞_ is fixed, we see that |*S*(*iω*)| stays large at higher frequencies rather than falling off quickly after its peak.

While any of these Tradeoffs could be studied in their own right, the important conceptual takeaway is that what underlies all of them is Bode’s integral theorem. In the same way that conservation laws provide a broad understanding the constraints on physic quantities (like momentum and energy), Equation (6) gives us a unifying framework for understanding the fundamental performance limitations of control systems. With this result in hand, we see that the performance tradeoffs shown here are simply different ways of tuning parameters to shape the function |*S*(*iω*)|. In the next section, we will apply some of these theoretical concepts to a particular biological circuit model. Though this model is more complex and nonlinear than those we have discussed so far, we will see that the same essential theoretical approach applies.

### 2.5 A Synthetic Growth Control Circuit

**Figure 4:**
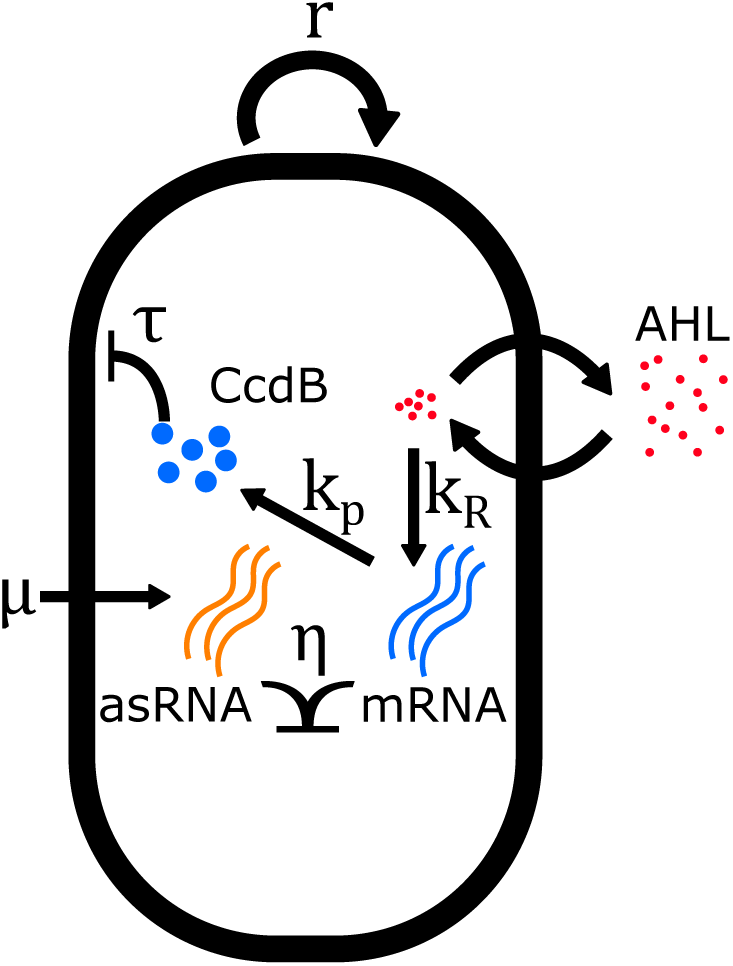
A Synthetic Growth Control Circuit. **A)** The circuit diagram for the dynamics described in equation (15). This circuit controls the growth of a bacterial population via the toxin CcdB. The concentration of CcdB is in turn regulated by a quorum sensing molecule AHL, whose mRNA can be sequestered by an antisense RNA [38]. This circuit is inspired by the work in [39] and has been implemented in [30]. This figure is adapted from [34].

Here we will use the results from previous sections to study a simple model of a synthetic sequestration feedback circuit based on the work in [30, 39], illustrated in figure 4A. The intended function of this circuit is to regulate the population level of a colony of *E. coli* via an external reference signal such as an inducer. We model the circuit with the following set of differential equations:

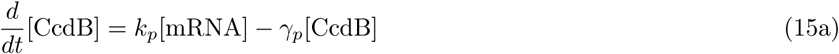

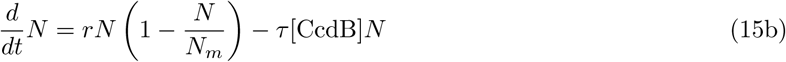

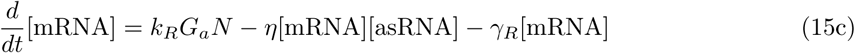

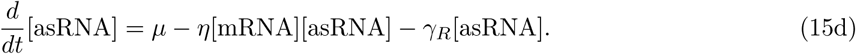

Quantities of the form [*·*] represent intracellular concentrations for each cell, and *N* represents the total number of cells. *N* follows logistic dynamics with an additional death rate due to toxicity *τ* proportional to the concentration of [CcdB] per cell. [CcdB] is a protein that is toxic to the cell, [mRNA] is the corresponding messenger RNA, the transcription of which we model as being induced by a quorum sensing ligand that is produced at a rate proportional to *N*, and [asRNA] is a short antisense RNA that has a complementary sequence to the CcdB mRNA, thus acting as a sequestering partner. The term *G_a_* = 10^−6^ nM captures the gain between *N* and mRNA induction mediated by the quorum-sensing molecule AHL.

As before, we will analyze a linearized version of this circuit. To do this we must first compute the steady-state values, shown in table 1. The linearized dynamics can now be written as

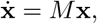

where

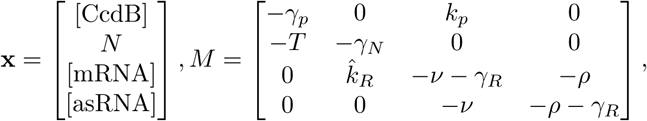

and 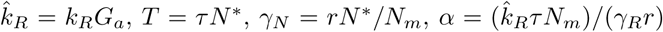, *v* = [asRNA]*, and *ρ* = [mRNA]*. From this, we can again derive stability results in the limit of large *η*. In terms of the parameters in *M*, we get a similar relationship to that of inequality (12), with the introduction of heterogeneous degradation rates:

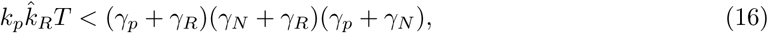

and the corresponding stability measure

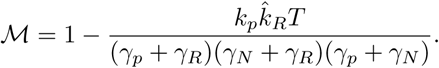

**Table 1:**
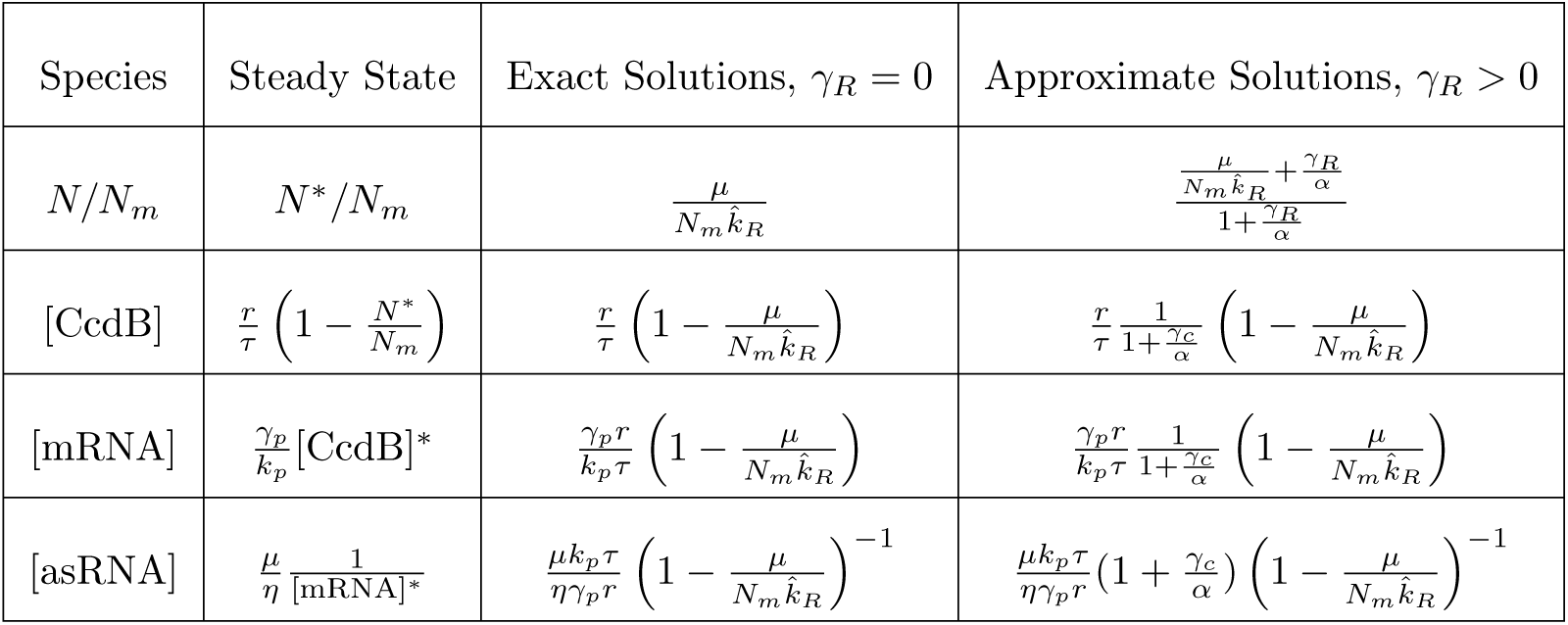
Steady state parameter values derive from equation (15). For the case *γ_R_* = 0 these solutions are exact, while they are approximated (assuming *η* large) for *γ_R_* > 0.

| Species | Steady State | Exact Solutions, $\gamma_R = 0$ | Approximate Solutions, $\gamma_R > 0$ |
| --- | --- | --- | --- |
| $N/N_m$ | $N^*/N_m$ | $\frac{\mu}{N_m \hat{k}_R}$ | $\frac{\frac{\mu}{N_m \hat{k}_R} + \frac{\gamma_R}{\alpha}}{1 + \frac{\gamma_R}{\alpha}}$ |
| [CcdB] | $\frac{r}{\tau} \left(1 - \frac{N^*}{N_m}\right)$ | $\frac{r}{\tau} \left(1 - \frac{\mu}{N_m \hat{k}_R}\right)$ | $\frac{r}{\tau} \frac{1}{1 + \frac{\gamma_c}{\alpha}} \left(1 - \frac{\mu}{N_m \hat{k}_R}\right)$ |
| [mRNA] | $\frac{\gamma_p}{k_p} [\text{CcdB}]^*$ | $\frac{\gamma_p r}{k_p \tau} \left(1 - \frac{\mu}{N_m \hat{k}_R}\right)$ | $\frac{\gamma_p r}{k_p \tau} \frac{1}{1 + \frac{\gamma_c}{\alpha}} \left(1 - \frac{\mu}{N_m \hat{k}_R}\right)$ |
| [asRNA] | $\frac{\mu}{\eta} \frac{1}{[\text{mRNA}]^*}$ | $\frac{\mu k_p \tau}{\eta \gamma_p r} \left(1 - \frac{\mu}{N_m \hat{k}_R}\right)^{-1}$ | $\frac{\mu k_p \tau}{\eta \gamma_p r} \left(1 + \frac{\gamma_c}{\alpha}\right) \left(1 - \frac{\mu}{N_m \hat{k}_R}\right)^{-1}$ |

A notable difference about this circuit is that stability is implicitly dependent on *μ*. This is because *μ* appears in *N*\*, which determines the values of *γ_N_* and *T*. Given that the function of this circuit is to control cell proliferation, it is natural to ask what steady-state levels of *N*\* are achievable for a given set of parameters. Because the scale of *N*\* is set by *N_m_*, we can non-dimensionalize the population size with the term *N*\*/*N_m_*. In the case *γ_R_* = 0, we can recast equation (16) as:

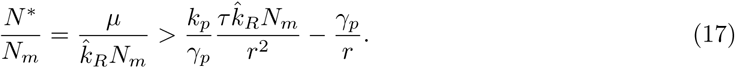

One immediate result of inequality (17) is that, if the following holds:

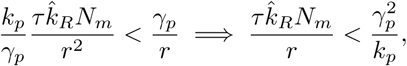

then the steady-state *N*\* is stable for any *μ* such that 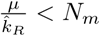 (the steady-state value of *N*\* cannot exceed the carrying capacity *N_m_* in equation (15b) from the nonlinear model). This constraint is also implicit in the steady-state value [asRNA]*, which is infinite if 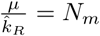. Because the right-hand side of the inequality has a factor of 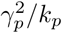, it is possible to improve performance without changing the steady-state concentration of [CcdB] by increasing both *k_p_* and *γ_p_* simultaneously, effectively increasing the protein’s turnover rate. If the right-hand side of inequality (17) is positive, then we see that the system’s performance is constrained, in that there is a certain population threshold below which *N*\* cannot be set. Just as in the previous section, as the system approaches this threshold it will become increasingly oscillatory. These effects were observed experimentally in [40], which uses the same general growth control architecture as in [39].

This section illustrates two key points, the first being that the general theoretical results from our initial analysis can be adapted to specific biological-motivated models of control. The second more general takeaway is that systems that look on the surface to be both biologically and mathematical distinct, e.g. a linear model of a chemical reaction network and a nonlinear population-level growth control circuit, have the same underlying structure. We often think of linearization simply as a method of approximation, but its real power often lies in showing us the connection between seemingly different mathematical models. In this case, it becomes clear what the analogous production and degradation rates are in equation (1) and equation (15).

This type of system-level theory allows us to abstract away details to see that seemingly different problems can be tackled with the same class of tools. In [34], we delve into simulations using biologically plausible parameter values and demonstrate that controller degradation can dramatically improve the circuit’s performance at relatively little cost.

### 2.6 Controlling Autocatalytic Processes

The general approach of the results presented so far has been to analyze in detail the simplest classes of networks that can be controlled by sequestration feedback. Going forward, it will be important to study networks where both the process and controller have more complex architecture. At the controller level, the sequestration mechanism alone only implements integral feedback. It will be useful to investigate mechanisms that could robustly implement proportional and derivative control mechanisms with the ultimate goal of synthesizing full proportional-integral-derivative (PID) controller [18, 41] in synthetic circuits.

It will also likely be essential to explore other mechanisms of implementing feedback control in living systems. Several mechanisms for biological control that are currently being explored include: paradoxical extracellular signaling inspired by process regulation [42] and post-translation mechanisms such as multi-protease regulation. Using control theoretical tools, it will be important to develop models for these biological controllers and assess their stability and performance. Researchers in bioengineering will likely benefit from having multiple mechanisms of feedback control to choose from, depending on the particular application.

Our results thus far has focused on the application of sequestration feedback to processes that are open-loop stable. It will likely be important to study the case of unstable processes, which can occur in autocatalytic networks such as the one involved in glycolysis and other metabolic processes. In control theory, unstable processes lead to a modified version of Bode’s integral theorem:

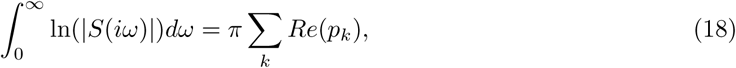

where *Re*(*p_k_*) is the real part of the unstable eigenvalues. Larger values of 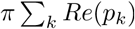 correspond to more global sensitivity to disturbances and harsher performance tradeoffs.

**Figure 5:**
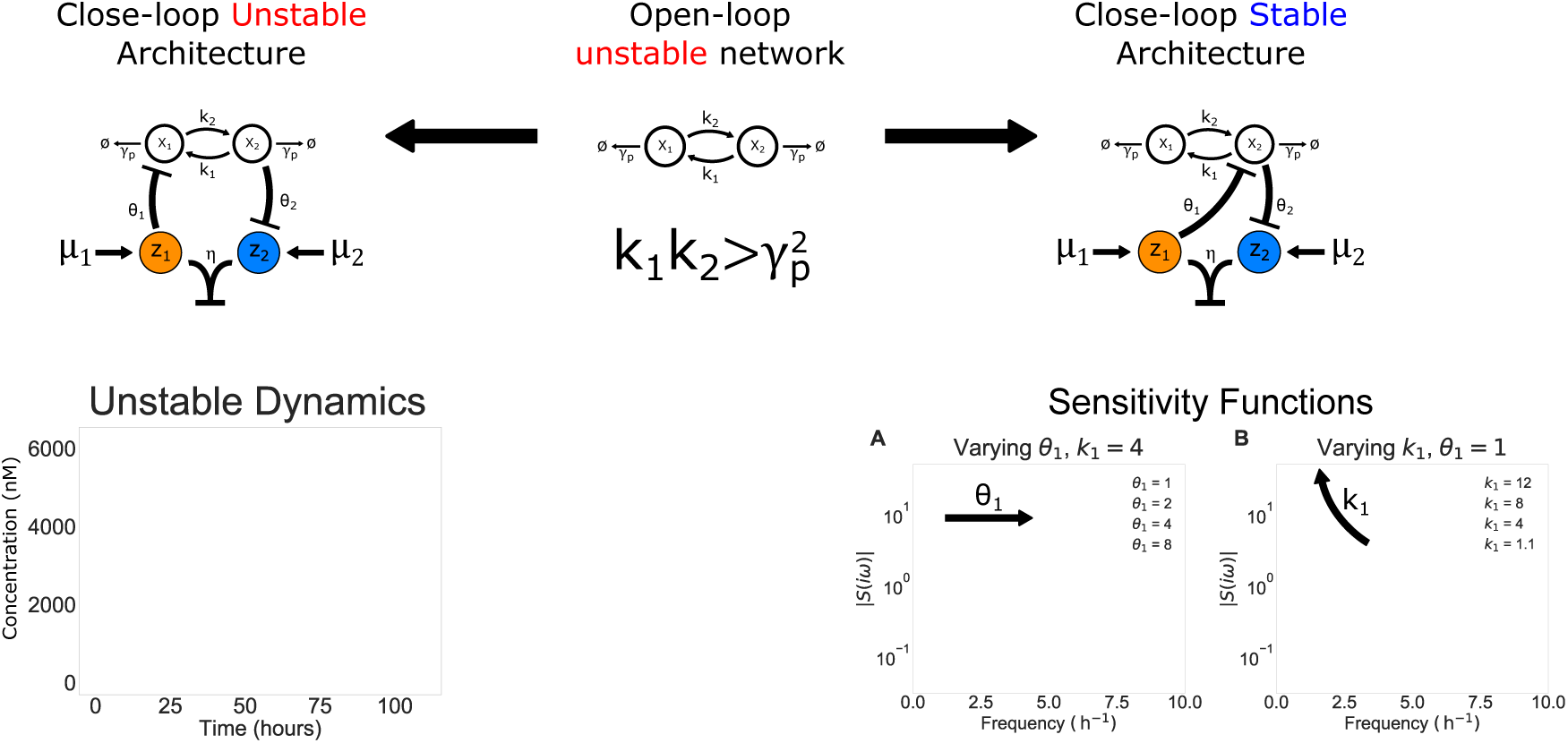
Using Sequestration to Control an Unstable Network. Here we take an unstable process (center) and study two different sequestration-based control architectures (left and right). This network is unstable so long as *k*_1_*k*_2_ > *γ*^2^. The repressive architecture on the left is *intrinsically* unstable, in that there are no values of the the control parameters that lead to the system reaching a stable steady state. A representative simulation of the unstable dynamics is presented below the architecture diagram. In contrast, the repressive architecture on the right is not only stabilizing, but *intrinsically* stabilizing. Any non-zero parameter values that result in positive steady-state concentrations of species will yield a stable closed-loop network. Panel **A** shows the sensitivity function as *θ*_1_ varies for a fixed value of *k*_1_ = 4 h^−1^. In this case, equation (6) tells us that the integrated area of the |*S*(*iω*)| will be constant as *θ*_1_ varies, because *θ*_1_ does not effect the location of unstable poles. In panel **B**, *θ*_1_ = 1 h^−1^ is fixed and *k*_1_ varies. This will change the location of the unstable pole, and we see a consequent change in integrated area of |*S*(*iω*)|, with large values of *k*_1_ leading to higher overall sensitivity of the system. In all simulations we take *θ*_2_ = *k*_2_ = *γ_p_* = 1 h^−1^, = 1000 h^−1^ nM^−1^, *μ*_1_ = 10 nM h^−1^, and *μ*_2_ = 110 nM h^−1^.

To demonstrate the nuance and complexity added by unstable processes, we demonstrate two seemingly similar control architectures that yield diametrically opposed behavior. As a simple model of an unstable process, we will use the process described in figure 5, which has the following dynamics:

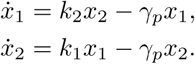

Since the system is linear, it is straightforward to check that the system is unstable when 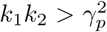. Because of the instability of the process, our controller will need to be repressive rather than activating, as it has been throughout the paper. The left panel of Figure 5 describes a plausible control architecture for such a system:

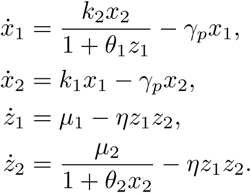

Here *z*_1_ represses *x*_1_ and *x*_2_ represses *z*_2_. Intuitively, if *x*_2_ is large then *z*_1_ will be reduced, increasing the amount of *z*_1_ which in turns reduces the amount of *x*_1_ and *x*_2_. We prove that this controller is actually *incapable* of stabilizing an unstable process, in that there are no parameters for which the closed-loop system is stable (section S4.4). If, however, we instead have *z*_1_ directly repress *x*_2_ (figure 5, right):

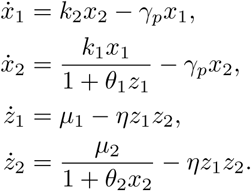

it is not only possible to stabilize the closed-loop system, but the system is *intrinsically* stable. So long as the system has positive parameter values and steady-state concentrations, we recover robust precise adaptation as presented in the earlier sections (section S4.4). While the stable process architecture could either be stable or unstable in closed-loop, this unstable process architecture confers a sort of inherent closed-loop stability that is quite surprising. If the system were linear, this would not be possible. Stability is a direct result of the nonlinearity introduced by repression. There is, however, a limitation: equation (18) tells us that a very unstable process 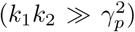 must exhibit extreme disturbance amplification. In terms of reference tracking, this implies that even the intrinsically stable controller will potentially have very bad transient behavior (e.g. extreme overshoot and ringing as the system stabilizes). While we can use the techniques developed in this paper to mathematically prove why these two architectures behave so differently, we have little biological insight into the architectural requirements for a stabilizing sequestration feedback controller. In the future we hope to develop a more general theoretical understanding of which architectures can confer stability to unstable networks.

## 3 Discussion

The development of synthetic biomolecular controllers could enable bioengineering to yield new solutions to problems in drug synthesis, immune system engineering, waste management and industrial fermentation [43–45]. In their current state, however, most current synthetic circuits lack the requisite robustness and scalability required of industrial technologies. The application of control theory to synthetic biological controllers aims to ensure that they function robustly in different host organisms and signaling contexts, despite perturbations from an uncertain environments.

The recent development of sequestration feedback controllers represents a promising step towards a general framework for implementing control in biological networks. This is best demonstrated by the rapid experimental progress towards implementing these controllers in a variety of contexts and with different sequestration mechanisms [28, 30, 46–48]. As these controllers become widely used, we believe that the theoretical results in this paper will not only provide a broad theoretical perspective on how the parameters of these networks interact to determine circuit performance, but also provide practical design rules that will tune circuit behavior in order to meet performance requirements. We begin to investigate these rules in [34], where we recast some of the results presented here (as well as some standalone results) as high-level architectural principles for understanding the performance of sequestration feedback circuits.

In the first half of the 20th century, the development of a cohesive theory of feedback control by Hendrik Bode, Harry Nyquist, and many other foundational thinkers facilitated the rapid development and proliferation of control systems in fields such as aerospace, electrical, and chemical engineering. The work presented here provides a link between the tools from classical control theory and contemporary problems in synthetic biology. In particular, we showed that it is possible to explicitly describe parametric conditions that determine stability, performance tradeoffs, and hard limits for a class of sequestration feedback controllers. While these limits can each be evaluated on their own, we observe that they can all be interpreted as different aspect of Bode’s integral theorem. This result acts like a fundamental conservation law for the performance of feedback control systems. By understanding these general theoretical constraints, we can gain a broad understanding of what is and is not achievable with a given control architecture.

## Author Contributions

Conceptualization and Methodology, NO, AAB, FX, YPL, JCD, and RMM; Formal Analysis, NO, AAB, FX, and YPL; Software, NO, AAB, and YPL; Writing, NO, AAB, and FX; Supervision and Funding, JCD and RMM.

## Acknowledgements

The authors would like to thank Reed McCardell for providing insight into the synthetic growth circuit, and Hana El-Samad for providing feedback on the manuscript. The project was sponsored by the Defense Advanced Research Projects Agency (Agreement HR0011-17-2-0008). The content of the information does not necessarily reflect the position or the policy of the Government, and no official endorsement should be inferred.

## Declaration of Interests

The authors declare no competing interests.

## S4 Supplemental Information

### S4.1 The Stability Criterion

We consider the mathematical model of the sequestration network described in equation (1). This mathematical model has a nonlinear term introduced by the sequestration dynamics. To evaluate its properties of stability and performance, we first linearize its dynamics. We can then describe the block structure of the linearized system in terms of the following matrices:

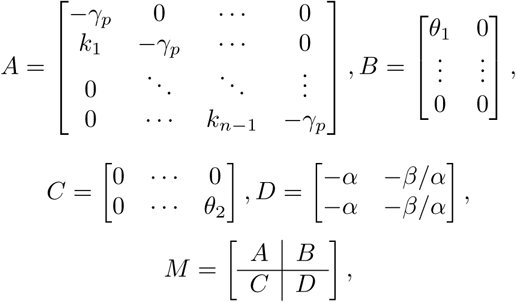

where 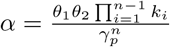 and *β* = *β*. The linearized dynamics will then be of the form

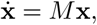

where

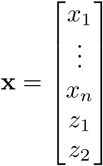

To prove our main stability result, we will analyze the characteristic polynomial of *M*, *p*(*s*). The roots of *p*(*s*) correspond to eigenvalues of *M*. In general it is difficult to analyze these roots, however we will see that the *p*(*s*) has a great deal of useful structure which we can exploit. First, we have to write down what *p*(*s*) actually is.

#### Lemma S1.

*The characteristic polynomial of M is*

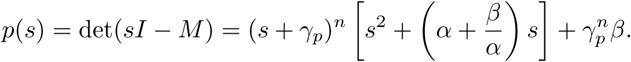

#### Proof.

We start by using the result that, for a block matrix such as *M*, we can use the classical result from linear algebra

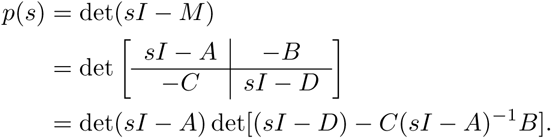

Since *A* is lower-triangular, we see immediately that the first term is

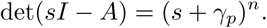

To analyze the second term, we first focus on computing *C*(*sI −A*)^−1^*B*. Because of the sparse structure of *B* and *C*, we have

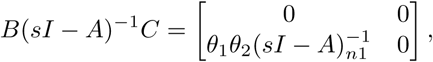

where 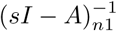 is the bottom-left most entry of (*sI − A*)^−1^. Using Cramer’s rule, we can compute

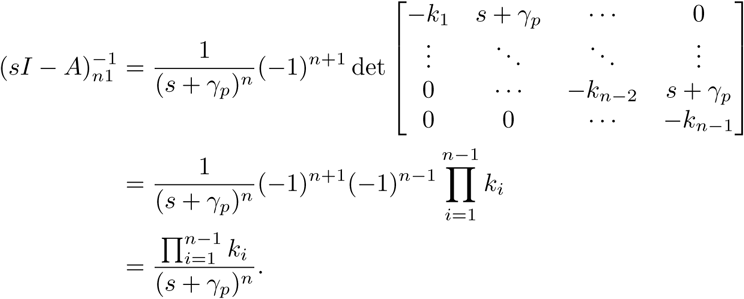

Combing these results, we see that

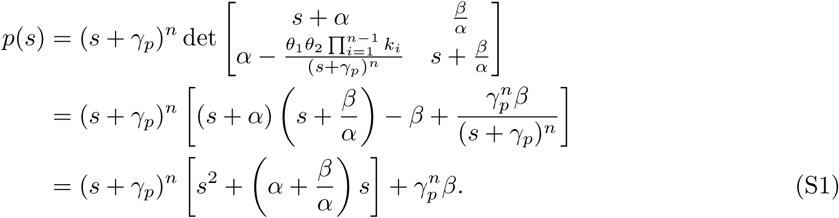

□

We can now use this result about *p*(*s*) to analyze the stability of the linearized sequestration feedback system.

#### Theorem S2 (Eigenvalue Classification Theorem)

*For a given n and β* ≫ *α*^2^,*αγ_p_, the eigenvalues λ of M has a parameter-independent classification of the form* 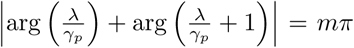 *for an integer m*.

#### Proof.

To study the eigenvalues of *M*, we will analyze the roots of *p*(*s*). We begin by by substituting *s* = *γ_p_σ* in equation (S1) and setting *p*(*σ*) = 0:

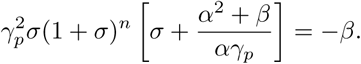

Taking the limit of strong feedback (*β* ≫ *α*^2^,*αγ_p_*), this equation reduces to

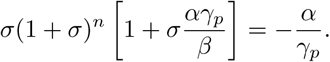

From this relationship we see that *p*(*σ*) has one large real root at 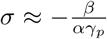. If we plug this into the phase constraint equation, this gives a phase of (*n* + 1)*π*. We will say the index of this root is *n* + 1. If 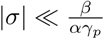 we get the simplified magnitude constraint

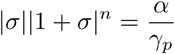

and the phase constraint

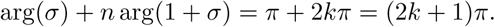

We can see from this that the maximum phase possible is *n* + 1 and that any each of the indices will be of the form 2*k* + 1 (i.e., odd integers). Because the magnitude constraint is independent of *k*, fundamentally we can have phase indices for any odd integer *m* such that |*m*| ≤ *n* + 1.

First we will see what conditions can produce purely real roots. If *σ* is real and *σ* > 0, then

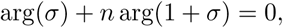

which violates the phase constraint. This implies that, if there are unstable roots, they are not purely real. If −1 < *σ* < 0, then

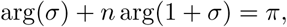

and we can have stable real roots with index 1. The magnitude constraint tells us that we will have a pair of these real roots if 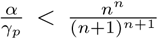 (which have index 1) with a bifurcation that generates conjugate pairs of roots when 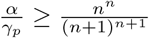. These conjugate roots will have indices ±1.

An immediate result of these observations is that, for any positive odd integer *m* such that 1 < *m* < *n* + 1, roots cannot be purely real and must come in conjugate pairs *±m*. If *n* is odd, then we will have a conjugate pair of roots for each *m* ∈ [3, *n* − 1], either a pair of small real roots or a conjugate pair for *m* = 1, and a single large negative real root for *m* = *n* + 1.

If *n* is odd, then the situation will be almost the same except for the fact that there will be a second real root with index *n* + 1. By some simple accounting, this analysis accounts for all *n* + 1 roots of *p*(*σ*), which correspond to roots of *p*(*s*) by a simple rescaling by 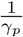.

□

#### Theorem S3 (Production-Degradation Inequality)

*Let M be the matrix corresponding to a linearization of the sequestration feedback system with two control molecules (z*_1_ *and z*_2_*) and n process species. In the limit of strong feedback* (*β* ≫ *α*^2^,*αγ_p_*), *the system is stable if and only if* 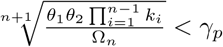 *where Ω_n_ is a constant that only depends on n. Further, when the system has purely imaginary eigenvalues the frequency of oscillation will be* 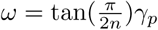.

#### Proof.

We will prove the results by finding parametric conditions that will result in purely imaginary eigenvalues, and then study what happens to the stability of the system when those parametric conditions do not hold (i.e. equalities become inequalities). To do this, we generalize a technique from [26], where we evaluate *p*(*s*) = 0 on the imaginary axis. In particular, we pick the change of variable *s* = *iω*\**γ_p_*, where *ω*\* is a positive real number (which we can assume without loss of generality because complex roots come in conjugate pairs), and evaluate *p*(*ω*\*). This yields the equations

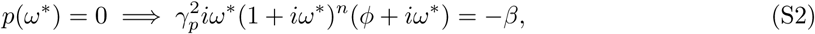

where 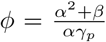. If we take the magnitude and phase of the the left-hand side of equation (S2), we get the constraints

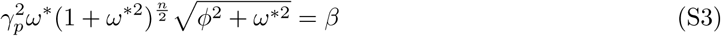

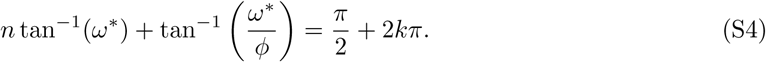

From theorem S2 we know that, in the limit of strong feedback, all complex eigenvalues have magnitude much less than *ϕ*, therefore tan^−1^(*ω*\*/*ϕ*) → 0. From these observations, we get the simplified relationship

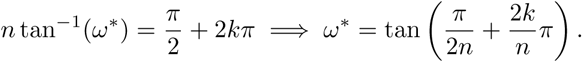

and equation (S3) becomes

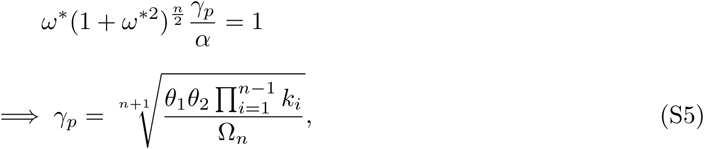

where 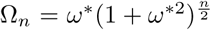. We can think of the parametric constrain equation (S5) as the boundary between stable and unstable behavior in the system. Because the left-hand side of equation (S3) is monotone in *ω*\*, we can infer that *ω*\* is unique and consequently there can only be one point in parameter space where there exist purely imaginary eigenvalues.

The final step is to study what happens when equation (S5) does not hold. First we look at the regime 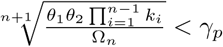. Again using the uniqueness of *ω*\*, if we understand the stability behavior of the system for a particular value of *γ_p_* in this regime, the same stability behavior must hold for all *γ_p_* in this range. Because of this, we can first examine the range where *γ_p_* is large. Intuitively, if degradation is sufficiently stronger than production then all species subject to degradation should converge to 0. To prove this rigorously, we will first search for roots with a large magnitude. If we apply the strong feedback limit to the characteristic equation from equation (S1), we get

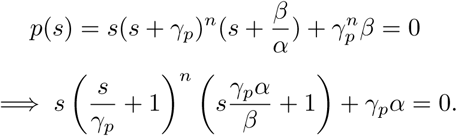

When |*s*| ≫*γ_p_α*, the characteristic equation will have the approximate form

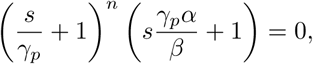

which gives us *n* roots at − *γ_p_* and one root at 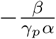. Since equation (S1) is order *n*+2, we know there is one remaining root outside of this regime. Next, we search for the final small root 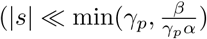, which gives relationship

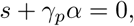

which gives a final small root at − *γ_p_α*. Since each of the *n* + 2 roots is negative, the system is stable for all 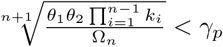.

Now we examine the regime 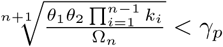. Here we will use a different technique, as taking the analogous limit of very small *γ_p_* is less straight-forward to analyze. To start, we will perform a change of variable *s* = *γ_p_σ*, where *σ* ∈ ℂ. We will again using the strong feedback limit, and study roots near the stability boundary, such that the characteristic equation still has the general form

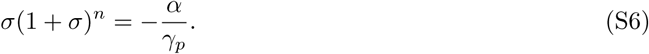

If we write *σ* = *a* + *ib*, we have the magnitude constraint

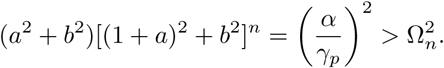

We also get the phase relationship

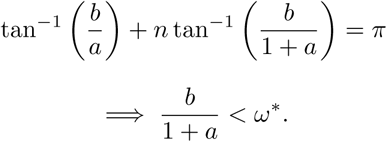

Combining these relationships, we get

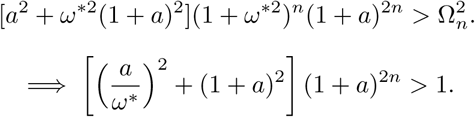

Since *a* = 0 at the stability boundary, there must be a regime of parameters sufficiently close to the boundary such that |*a*| ≪ *ω*\*, for which we have the relationship

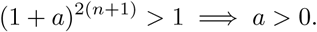

This proves the existence of an unstable point when 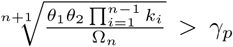, which implies that all parameters in this regime will yield unstable dynamics (so long as the strong feedback assumption still holds).

□

We note that, though previous results studied the regime of strong feedback (*β* large), the core assumption that was made is that the quantity

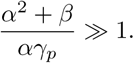

We note that there is an entirely different way to achieve this, by making *α*^2^ ≫*β*,α *γ_p_*. In this regime, all of the previous results follow in almost exactly the same way, except for changes to the constants involved. It is relatively straightforward to show that the characteristic equation for the system reduces to

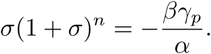

Following the same steps from the previous proofs, we can find that instability now occurs when

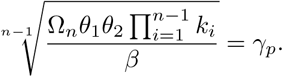

Interestingly, the stable regime is now

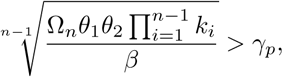

the opposite of what occurs in the strong feedback limit. One interpretation of these results as a whole is that stability is achievable when either controller sequestration or process degradation are individually large, but not when both are large simultaneously.

### S4.2 The Sensitivity Function

The sensitivity function *S*(*s*), *s* ∈ ℂ is the transfer function between an input reference to a system and output error [18]. It is particularly useful to examine |*S*(*iω*)|, which corresponds to the magnitude of *S* given a purely oscillatory disturbance. If |*S*(*iω*)| > 1, then the system will amplify disturbances at a frequency *ω*. Conversely, if |*S*(*iω*)| < 1 then the system will attenuate disturbances at frequency *ω*.

Define *P* (*s*) and *C*(*s*) to be the transfer function between inputs and outputs of the process and controller, respectively. It is a standard result in control theory that

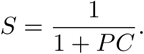

In general, for a linear system

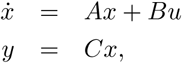

the transfer function has the form *H*(*s*) = *C*(*sI − A*)^−1^*B*. For the sequestration feedback system, we have that

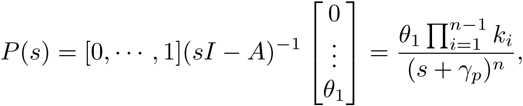

where just as before we use

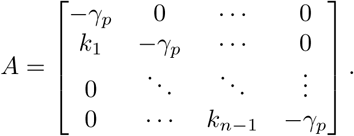

Similarly, we have that

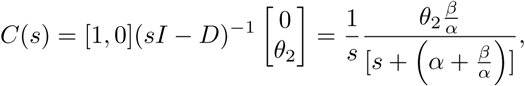

where

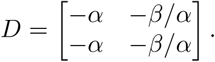

Note that *C*(*s*) has a factor if 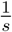, indicating that it corresponds to an integrator. From *P* and *C*, we see that

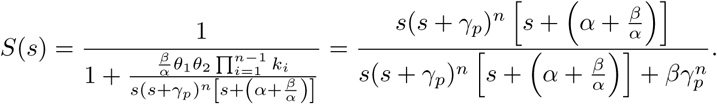

If we again take the limit 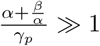 and substitute *s* = *γ_p_σ* we get the approximation

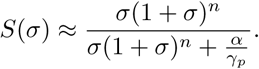

Ideally we would like to analyze ∥*S*(*iω*)∥_∞_ = max*_ω_*|*S*(*iω*)|, however this is difficult to compute in general. A lower bound for this term can, however, be easily computed by evaluating a particular value of *ω* close to the maximum. Specifically, we will use 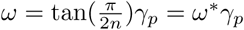. At *σ* = *iω*\*, we get

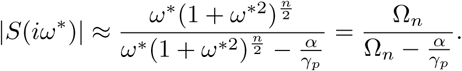

From our previous results, we know that the system is purely oscillatory when 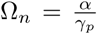, which corresponds to |*S*(*iω*\*)| = ∥*S*(*iω*)∥_∞_ = ∞. This gives the intuitive result that the system is infinitely sensitive to a periodic disturbance at *ω* = *ω*\**γ_p_* when 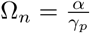. In general, we will have that

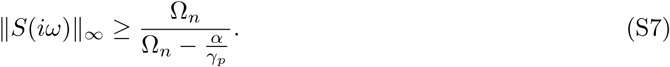

For the special case of *n* = 2, we can explicitly derive an even tighter bound than the one in inequality (S7). First, we can explicitly compute

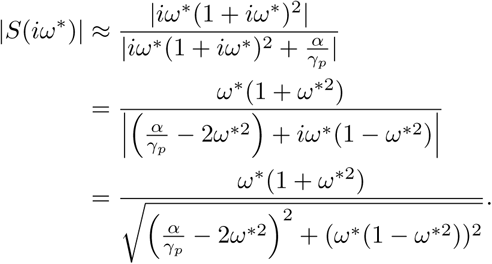

Much of the complexity in this equation comes from the denominator, which can be simplified if we pick *ω*\* such that either the real or imaginary part is 0. If we plug in *ω*\* = tan(*π*/4) = 1, the complex part of the denominator vanishes and we recover the original bound:

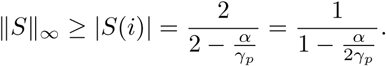

To set the real part to zero, it must be the case that

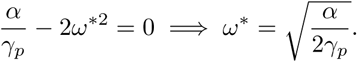

Plugging this in, we get that

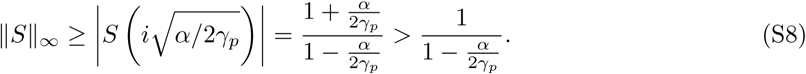

We see that this new bound is strictly greater than the one derived in inequality (S7), and therefore is a better approximation of ∥*S*∥_∞_. While inequality (S7) generalizes to all value of *n*, the latter bound unfortunately requires us to find real roots of order *n* polynomials, which scales poorly for this problem.

### S4.3 Sequestration Feedback with Controller Species Degradation

#### S4.3.1 Steady state analysis

Following the same notation as the previous sections, we can model the role of controller degradation as

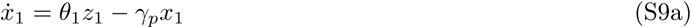

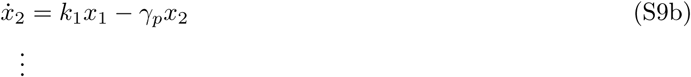

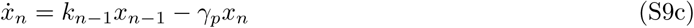

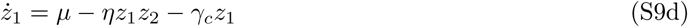

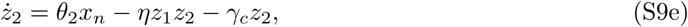

where *γ_p_* is the degradation of the process species *x_i_* and *γ_c_* is the degradation rate of the control species *z*_1_ and *z*_2_. At a high level we will proceed much in the same way as we did previously, however we will see that nonzero controller degradation leads to several technical challenges that do no appear when *γ_c_* = 0. The first of theses arises from simply solving for the steady values around which we will linearize the model. Where previously we used the fact that 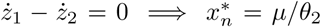, where * denotes a steady-state value. to subsequently solve for all other steady-state concentrations, we are now left with the messier relationship

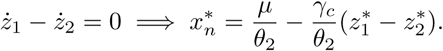

This implies that, for *γ_c_* > 0, we expect *x_n_* to differ from the desired steady-state *μ/θ* by some error that depends on the values of 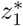 and 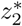. Since this error is almost surely a function of many other parameters, we essentially lose the robust precise adaptation property where 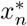 is completely independent of the network’s parameters. We will first calculate a general form for 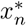, then derive a large *η* limit thats make further calculations tractable.

To begin, we use equations (S9d) and (S9e) to derive the relationships

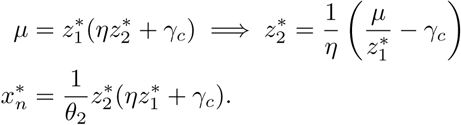

Combining these equations, we find that

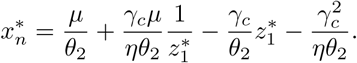

Finally, we observe that

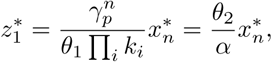

which yields the relationship

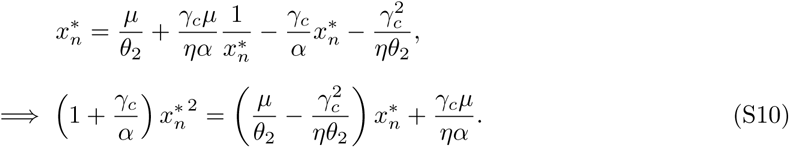

While this quadratic can be solved explicitly, the result can be greatly simplified by again taking the limit of large *η*. Here the sense in which we take this limit is such that 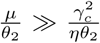 and 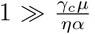. These reduce to the condition

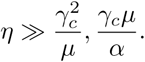

Combined with the previous assumption about the size of we now have a large number of conditions to fulfill, however we find that in practice we rarely are in parameter regimes where a great deal of tuning needs to be done to satisfy everything. That being said, we can use this limit to reduce equation (S10) to

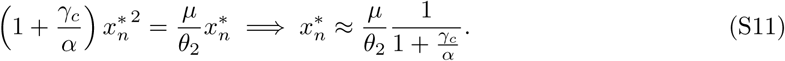

Using the same approximation, we can also compute

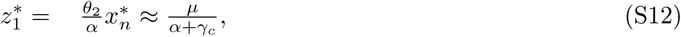

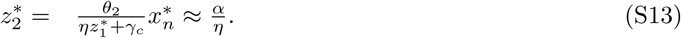

These will be useful for computing the linearized dynamics of the system in the next section.

As a sanity check, we can immediately see that 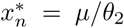 when *γ_c_* = 0, as expected. For *γ_c_* > 0, equation (S11) captures the steady-state error relative to the set point *μ/θ* induced by nonzero controller degradation. We see that, so long as the ratio *γ_c_/α* ≪ 1, error will be negligible. What is unclear at this point is under what conditions this can be achieved while still ensuring stability of the overall system. To this end, we will now characterize stability and performance for *γ_c_* > 0.

#### S4.3.2 Linearized dynamics and characteristic equation

Here we present results analogous to those in section S4.1, omitting detailed proofs since the structure of the argument from this point on is essentially identical to what was show in the previous section. Because the only nonlinear terms in our system are in equations (S9d) and (S9e), the only matrix to change in our linearization from section S4.1 is

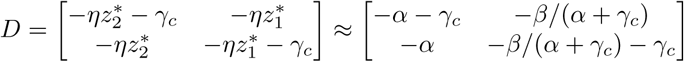

Using this *D* matrix and proceeding with precisely the same calculation as before, we can derive the characteristic equation for the system:

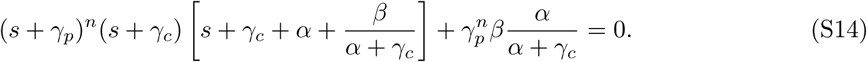

We again take the appropriate limit of *β* ≫ (*γ_c_* + *α*) *γ_p_*, (*γ_c_* + *α*)^2^ to follow the same argument as in section S4.1 to get the simplified expression in terms of *σ* = *s/γ_p_*:

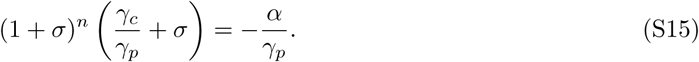

First we note that, when *γ_c_* = 0, we recover equation (S6) as expected. Proceeding as before, we can write the characteristic polynomial in terms of phase and magnitude constraints for *σ* = *iω*\*:

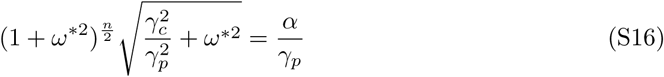

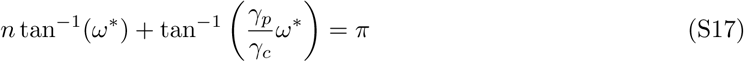

#### S4.3.3 Stability analysis

Unfortunately, the additional complexity in equation (S17) makes it challenging to write down the sort of explicit closed-form expressions for stability seen in theorem S3. While we can write out explicit stability conditions for *n* = 2, we will need to study particular parameter regimes for *n* > 2 as the summation relationship for tan^−1^ scales poorly.

To solve for *ω*\* in equation (S17) we make use of the inverse trigonometric identity

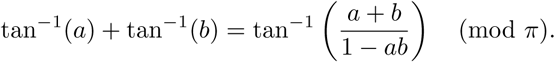

Applying this identity twice yields the relationship

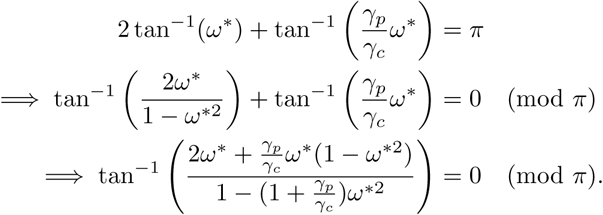

Since the only value for which tan^−1^(*x*) = 0 is *x* = 0, the problem reduces to solving the equation

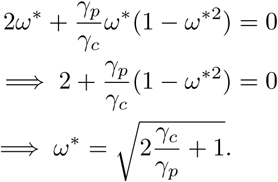

Combining this with equation (S16) yields the stability criterion

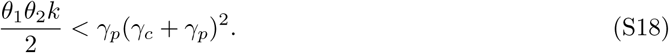

If we assume that we have full freedom to set control parameters, then inequality (S18) that it is possible to make the production rates *θ*_1_ and *θ*_2_ large, so long as there is a compensatory increase in *γ_c_*. This implies that we can, in a sense, sidestep the performance tradeoffs between speed and stability if we are willing pay a price in terms of *efficiency*, measured by the turnover rates of *z*_1_ and *z*_2_.

Next we will study what happens when *n* > 2. We note that there is an interesting topological distinction going from *n* = 2 to *n* > 2 which yields qualitatively different stability results. To see why this is the case, we return to equation (S17):

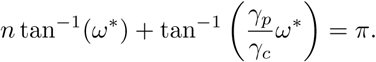

Recall that tan^−1^(*x*) < *π*/2 for all *x*. Because this is the case, when *n* = 2 it is always the case that 2 tan^−1^(*ω*\*) < *π*, implying that satisfying the phase condition is strictly contingent of the value of the term tan^−1^((*γ_p_/γ_c_*)*ω*\*). On the other hand, for *n* > 2, there exist values of *ω*\* such that *n* tan^−1^(*ω*\*) ≥ *π*, so depending on the relative magnitude of the ratio *γ_p_/γ_c_* satisfying the phase condition may or may not depend strongly on *γ_c_*.

If we look again at equation (S15):

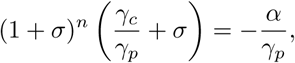

we notice that the only place in which *γ_c_* appears is in the ratio *γ_c_/γ_p_*. One natural approach to studying the solutions to this equation is to examine what happens at various limits, namely *γ_c_* ≪ *γ_p_*, *γ_c_* = *γ_p_*, and *γ_c_* ≫*γ_p_*. Here we will present results without going into formal detail, however the analysis can be made rigorous by analyzing equation (S17).

##### Case I

*γ_c_* ≪ *γ_p_*: This case is fairly straightforward, as it is it reduces to the case of no controller degradation. We recover the characteristic polynomial

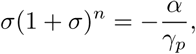

which has the same exact stability condition as in theorem S3.

##### Case II

*γ_c_* = *γ_p_*: This case is representative of what happens when controller and process degradation have the same order of magnitude. We use *γ_p_/γ_c_* = 1 in equation (S17) to find that the stability boundary is characterized by:

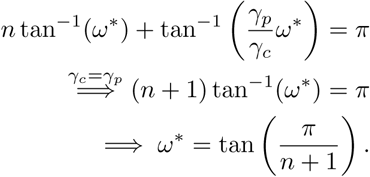

Here it is useful to define the quantity

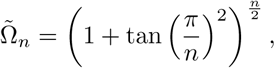

where 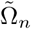 differs from the previously defined Ω*_n_* by a factor of 1/2 in the argument of the tangent term. Using this expression, we can use equation (S16) to derive the stability criterion

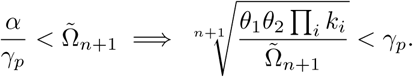

This condition is qualitatively the same as the one in theorem S3 up to a constant difference accounted for by the 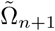 term.

##### Case III

*γ_c_* ≫ *γ_p_*: Following a similar line of reasoning as in the previous case, taking the limit *γ_p_/γ_c_* ≪1 in equation (S17) to show that

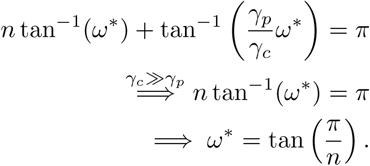

We again use equation (S16) to find that the stability boundary is set by the following relationship:

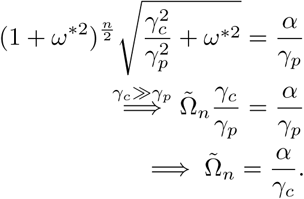

This implies that the stability criterion for this case is

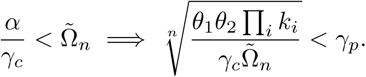

Notice that in the *n* = 2 case, *ω*\* is a function of *γ_c_* and the subsequent stability criterion depends on the term (*γ_p_*+*γ_c_*)^2^. This is quite different from the *n* > 2 cases where in each regime, the *γ_c_* dependence in *ω*\* disappears. Similarly, in the stability criterion we see a linear (rather than quadratic) dependence on *γ_c_*. This is a direct result of the previously mentioned topological difference between the *n* = 2 and *n* > 2 cases.

One interesting side effect of this results is that, when the system is purely oscillatory (on the stability boundary), the frequencies of oscillation may be dramatically different depending on *n*. Consider the case where *γ_c_* ≫*γ_p_* If *n* = 2, this frequency will be

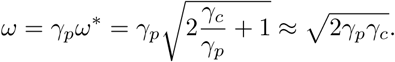

If *n* > 2, we use the results from Case III above to find

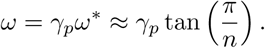

In the former case, *ω* scales with 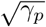, whereas in the latter case *ω* is independent of *γ_c_*. This implies that for large controller degradation rates we would expect much faster oscillatory modes for *n* = 2 than for *n* > 2.

#### S4.3.4 The effects of degradation on sensitivity and performance

Just as in section S4.2, we can write the generic sensitivity function for the linearized sequestration feedback system with degradation in terms of the variable *σ* = *γ_p_s* as

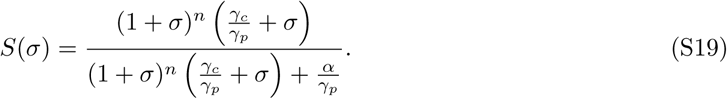

For the case *n* = 2, we can again derive an explicit lower bound for ∥*S*(*iω*\*) ∥_∞_:

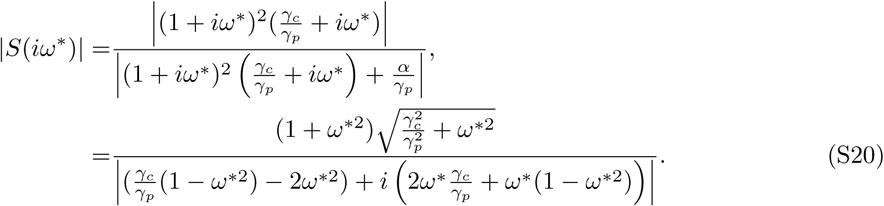

We can again solve for *ω*\* such that the real part of the denominator is zero:

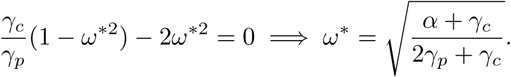

If we evaluate equation (S20) at *ω*\*, we can write the bound

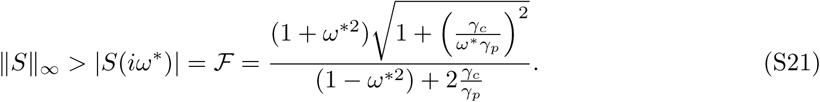

It is easy to check that, for *γ_c_* = 0, we recover the bound from inequality (S8). As *ω*\* approaches 1 + 2*γ_c_/γ_p_*, ∥*S*∥_∞_ will asymptotically increase to ∞. Alternatively, increasing *γ_c_* will decrease sensitivity, and consequently improve robustness. We can think of *γ_c_* as capturing the inefficiency of our controller (higher degradation means the control species are degraded before being used in a sequestration reaction). In these terms, we see that increasing *γ_c_* will reduce ℱ at the cost of increased steady-state error (see figure 3D-F). If we hold *ε* constant by varying both *γ_c_* and *θ*_1_, the we can decrease ℱ at the cost of on decreasing efficiency of the controller (see figure 3G-I). Finally, we can vary *γ_c_* and *θ*_1_ such that ℱ is constant, which leads to a tradeoff between steady-state error and efficiency (see figure 3J-L).

### S4.4 Controlling an Unstable Process

In all prior sections, we have assumed that the underlying process being controlled is open-loop stable. Here we will examine a simple model of an open-loop unstable process and describe which control architectures are capable of stabilizing the closed-loop system. To start, we will use a simple linear system as our process:

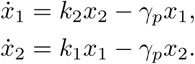

This system will be unstable when at least one eigenvalue of the matrix:

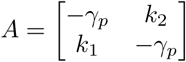

has positive real part. With some straightforward linear algebra we can find that the eigenvalues of *A* are,

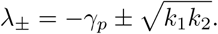

Because *k*_1_, *k*_2_, *γ_p_* > 0, we know that *λ*_−_ < 0 for all parameters. *λ*_+_, however, can be either positive or negative. In particular,

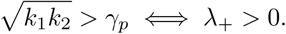

To facilitate our study of unstable processes, we will assume 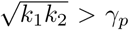 for the rest of the section. One immediate difference is that, due to the unstable process, any controller must now be repressive.

To model this, we will first study the following architecture (described in equation (19) and the left panels of figure 5):

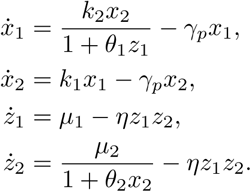

If *θ*_1_ = *θ*_2_ = 0, then this architecture reduces to the open-loop system described above. The controller topology is essentially the same as in the stable case, with the core difference that *z*_1_ represses *x*_1_ and *x*_2_ represses *z*_2_, where before these interactions were activating. Since now there is no reaction synthesizing *z*_2_, we must add in some external production rate *μ*_2_. We will again proceed by solving for the steady-state concentrations of each species and linearizing around these values. The steady-state concentrations are as follows:

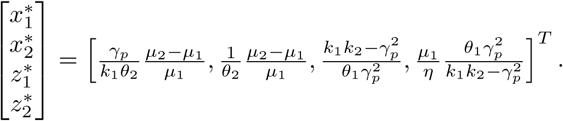

If we now linearize around this fixed point, we can define a new set of parameters:

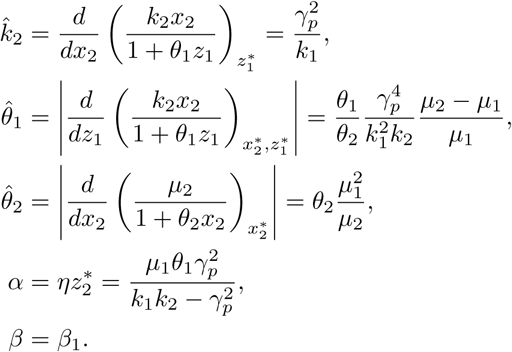

which characterize the linearized set of dynamics:

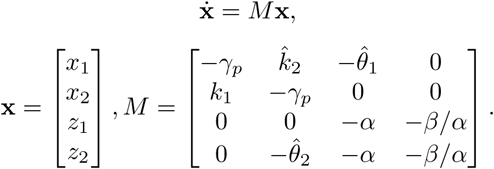

Following the same methods in section S4.1, we can derive the characteristic polynomial for *M*,

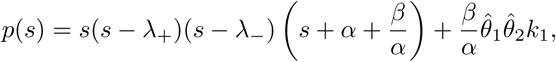

where we now use 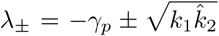. If we plug in 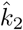, we see that *λ*_+_ = 0 and *λ*_−_ = −2*γ_p_*. The fact that the process’s eigenvalues change when comparing the open- and closed-loop systems is a byproduct of the fact that our original model was nonlinear, and is something that would not occur for a strictly linear system.

Again taking the limit of strong feedback, which here takes the form *β* ≫ *α*^2^, 2*αγ_p_*, and setting *p*(*s*) = 0, we get the equation

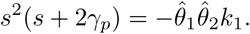

The corresponding phase constraint for this equation when *s* = *iω* is (after some algebra):

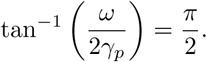

Since tan^−1^(*x*) < *π*/2, there are no parameter values for which this phase constraint is achieved.

Next, we consider an alternative architecture, shown in figure 5 and equation (20). This system is described by the same dynamics as before, except we now have *z*_1_ directly repressing *x*_2_ (rather than indirectly doing so via *x*_1_):

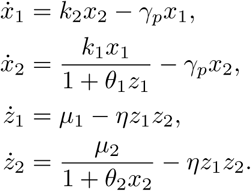

The steady-state values are almost identical to those of section S4.4, except we now have that

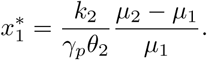

We can define another set of linearized parameters,

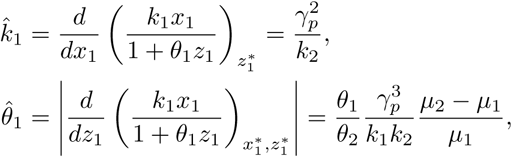

with 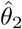, *α* and *β* the same as before. Our linearized dynamics are now described by the matrix

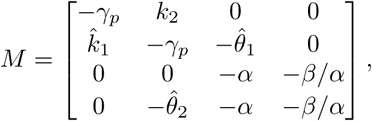

with a corresponding characteristic polynomial

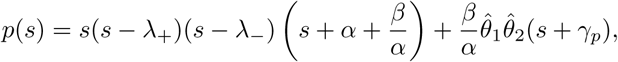

with 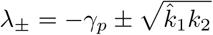. The limiting form of the characteristic equation is now

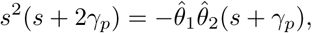

with the phase constraint

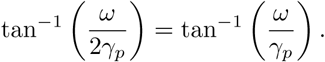

Unlike in the previous architecture, this constraint *is* achievable for *ω* = 0. this leads to a stability criterion of the form

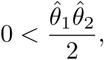

which implies that the linear system is intrinsically stable so long as the parameters are set such that the system has a positive steady-state concentrations (*μ*_2_ > *μ*_1_, 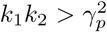). The in turn implies that the nonlinear system will be locally stable near the fixed point *independent* of the model’s parameters.

Finally, we can find the sensitivity function for the stabilizing architecture. This is somewhat complicated by the fact that the process transfer function varies with control parameters, so it is difficult to separate the process and the controller transfer functions. However we can use a convenient form

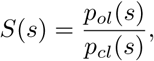

where *p_cl_*(*s*) is the characteristic equations for the closed-loop systems, and 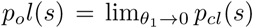 [8, 49]. Using

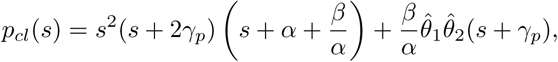

we get the sensitivity function (assuming large *β*):

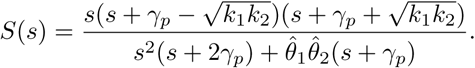

Note that it is important the we take care with the limits, as the roots of *p_ol_*(*s*) should reflect the eigenvalues of the unstable open-loop system. This is used to generate the right-hand plots in figure 5A and B.

